# Growth Mode and Carbon Source Impact the Surfaceome Dynamics of *Lactobacillus rhamnosus* GG

**DOI:** 10.1101/579953

**Authors:** Kirsi Savijoki, Tuula A. Nyman, Veera Kainulainen, Ilkka Miettinen, Pia Siljamäki, Adyary Fallarero, Jouko Sandholm, Reetta Satokari, Pekka Varmanen

## Abstract

Bacterial biofilms have clear implications in disease and in food applications involving probiotics. Here, we show that switching the carbohydrate source from glucose to fructose increased the biofilm formation and the total surface-antigenicity of a well-known probiotic, *Lactobacillus rhamnosus* GG. Surfaceomes (all cell surface-associated proteins) of GG cells grown with glucose and fructose in planktonic and biofilm cultures were identified and compared, which indicated carbohydrate source-dependent variations, especially during biofilm growth. The most distinctive differences under these conditions were detected with several surface adhesins (e.g., MBF, SpaC/A pilus and penicillin-binding proteins), enzymes (glycoside hydrolases, PrsA, PrtP, PrtR and HtrA) and moonlighting proteins (glycolytic, transcription/translation and stress-associated proteins, r-proteins, tRNA synthetases, Clp family proteins, PepC, PepN and PepA). The abundance of several known adhesins and novel moonlighters, including enzymes acting on casein-derived peptides (ClpP, PepC and PepN), increased in the biofilm cells grown on fructose, from which the surface-associated aminopeptidase activity mediated by PepC and PepN was further confirmed by an enzymatic assay. The classical surface adhesins were predicted to be more abundant on planktonic cells growing either on fructose (MBF and SpaA) or glucose (SpaC). An additional indirect ELISA indicated both growth mode- and carbohydrate-dependent differences in abundance of SpaC, whereas the overall adherence of GG assessed with porcine mucus indicated that the carbon source and the growth mode affected mucus adhesion. The adherence of GG cells to mucus was almost completely inhibited by anti-SpaC antibodies regardless of growth mode and/or carbohydrate source, indicating the key role of the SpaCBA pilus in adherence under the tested conditions. Altogether, our results suggest that carbon source and growth mode coordinate classical and nonclassical protein export in GG, which ensures the presence of an integral and coordinated system that contributes to resistance, nutrient acquisition and cell-cell interactions under different conditions. In conclusion, the present study shows that different growth regimes and conditions can have a profound impact on the adherent and antigenic features of GG, thereby providing new information on how to gain additional benefits from this probiotic.

## Introduction

Probiotic bacteria, applied in various aspects of the food and pharmaceutical industries (1, 2), exploit sophisticated strategies to enable the utilization of many nutrients, maintain viability and adherence to mucosal surfaces, and modulate host immunity in a beneficial way during industrial processes and upon their consumption (3). Unlike free-living cells, the biofilm mode of growth has been shown to improve existing probiotic features (antimicrobial and anti-inflammatory functions) in some *Lactobacillus* strains (4–6). In addition, switching from the planktonic to the biofilm mode of growth can provide bacteria, including probiotics, with means to overcome lethal conditions, including antimicrobial treatments, host immune defenses, and changes in pH, salt, temperature and nutrients (7). Biofilms comprise cells enclosed in an extracellular matrix (consisting of nucleic acids, exopolysaccharides (EPS), lipids and/or proteins) and typically display adherent growth on either abiotic or natural surfaces (8). Probiotics must be consumed at least daily to obtain most of their benefits, and new probiotic supplements prepared as biofilms, e.g., in biocompatible microspheres or encapsulated in microcapsules, could offer longer lasting probiotic effects compared to those prepared from free-grown cells (9–11).

One of the most documented and utilized probiotic *Lactobacillus* strains, *L. rhamnosus* GG, is known to adhere to the human intestinal mucosa via pilus (encoded by *spaCBA*) extending from its cell surface and to persist for more than a week in the gastrointestinal tract (GIT) of healthy adults (12). EPS is another factor that plays an important role as a protective shield against host innate defense molecules to improve adaptation in the GIT (13). However, reduced EPS synthesis has been linked to the increased adherence and biofilm formation of GG, which by uncovering specific adhesins necessary for biofilm formation, enabled the enhanced biofilm growth of this strain (14). The SpaCBA-pilus adhesin, which mediates the direct interaction with the host and abiotic surfaces, and MabA (the modulator of adhesion and biofilm), which strengthens the biofilm structure, are considered the central factors that mediate the biofilm formation of GG *in vitro* (15, 16). While the presence of biofilms in the colonic microbiome has been reported in several studies, GIT-associated probiotic biofilms have been found in specific niches of certain animal hosts (17, 18). GG can grow as a biofilm on inert substrates (15, 19, 20) and integrate well in a multispecies biofilm model with the potential to inhibit the growth of some cariogenic species (21). However, systematic studies aiming to uncover all surface-bound proteins on GG biofilms have not been conducted.

We previously demonstrated that the biofilm formation of GG in MRS medium under microaerophilic and anaerobic (5% CO_2_) conditions was protein-mediated (19), indicating that cell surface-associated proteins most likely affected the biofilm formation of this strain. In a multispecies biofilm model, GG was shown to reach high viable cell numbers with glucose or sucrose as the carbon source (22). Glucose and fructose can enhance the survival of GG in simulated gastric juice at pH 2.0 (23), while fructose also contributes to the *in vitro* adherence and antimicrobial activity of this probiotic (24, 25). These simple carbohydrates were recently found to prevent the colonization of a gut commensal bacterium, *Bacteroides thetaiotaomicron*, in the gut, an effect that was not observed with prebiotic fructo-oligosaccharides (26).

The present study aimed to uncover the effects of two simple carbohydrates, glucose and fructose, on the total surface-associated antigenicity of GG growing in planktonic and biofilm states. To explore these findings further, the surfaceome compositions from the same cells were compared, and some industrially relevant features were verified with phenotypic analyses. To the best of our knowledge, this is the first systematic study exploring surfaceomes of probiotic biofilms and demonstrating the importance of growth mode and carbon source in coordinating the adherent and industrial features of GG.

## Materials and Methods

### Culture media

*Lactobacillus rhamnosus* GG was routinely grown on commercial MRS agar (BD, Franklin Lakes, US) prior to propagation in modified MRS broth for planktonic or biofilm cultivations. The composition of this modified MRS was as follows: 10 g/L tryptone pancreatic digest of casein, 10 g/L beef extract powder, 5 g/L Bacto^TM^ yeast extract, 1 g/L Tween 80, 2 g/L ammonium citrate tribasic, 5 g/L sodium acetate, 0.1 g/L magnesium sulfate heptahydrate, 0.05 g/L manganese II sulfate monohydrate and 2 g/L dipotassium hydrogen phosphate trihydrate (pH 6.5-7). When appropriate, MRS was supplemented with 2% glucose (glc) (MRS-G) or 2% fructose (frc) (MRS-F) (Sigma-Aldrich, Espoo Finland) or less (1.75%, 1.5%, 1.25%, 1.0%, 0.75%, 0.5%, 0.25%, 0.1, or 0.05%).

### Biofilm formation and quantification

GG colonies from MRS agar plates were suspended in MRS supplemented with glc or frc at desired concentrations to obtain an OD_600_ = 0.15-0.2, from which 200 µL was removed and added to flat-bottomed 96-well plates (FALCON; Tissue Culture Treated, polystyrene, Becton Dickinson), and the plates were incubated at 37 °C under anaerobic conditions (5% CO_2_) for 24 h, 48 h or 72 h. Biofilm formation efficiency was assessed with crystal violet staining essentially as previously described (27). Briefly, the non-adherent cells were removed from the wells, and the biofilm cells were washed twice with deionized H_2_O. Adherent cells were stained with 200 μL of the crystal violet solution (0.1%, w/v) (Sigma–Aldrich, Munich, Germany) for 30 min at RT. Excess stain was removed by washing the cells twice with deionized H_2_O, and the stained cells were suspended in 200 μL of 30% acetic acid by shaking at RT (400 r.p.m. for 30 min). The density of the biofilms was recorded at 540 nm using an ELISA reader (Labsystems Multiskan EX). Biofilm experiments were performed several times, using at least eight technical replicates.

### LIVE/DEAD staining of biofilms and confocal microscopy

GG cells were suspended in MRS-G (2% glc) or MRS-F (2% frc) to obtain OD_600_ ∼ 0.15-0.2, from which 4 mL was added to uncoated 35 mm glass bottom dishes (MatTek, Ashland, MA, USA). Dishes were incubated at 37 °C under anaerobic conditions (5% CO_2_) for 48 hours for biofilm formation. Non-adherent cells were removed, and adherent cells were subjected to LIVE/DEAD viability staining using 5 µM of Syto9 and 30 µM of propidium iodide (PI) according to the manufacturer’s instructions (LIVE/DEAD^®^ *Bac*Light™, Molecular Probes, Life Technologies). Fluorescence images were captured with Zeiss LSM510 META confocal microscope using Zeiss LSM 3.2 software (Zeiss GmbH, Oberkochen, Germany). All biofilms were analyzed in duplicates.

### 1-DE immunoblotting

Protein samples from planktonic and biofilm cell surfaces for 1-DE (SDS polyacrylamide gel electrophoresis) and immunoblotting were prepared as follows. GG cells were suspended in MRS-G or MRS-F to obtain an OD_600_ = 0.15-0.2. The cell suspensions were split in two; half (2.5 mL) of the suspension was cultured under planktonic conditions (Falcon tubes) and half (2.5 mL/ tube) was cultured under biofilm-formation conditions (2.5 mL per well in a flat-bottomed 24-well plate; FALCON - Tissue Culture Treated, Polystyrene, Becton Dickinson) at 37 °C in the presence of 5% CO_2_. The cell densities of planktonic cultures and biofilms, after suspending the adherent cells in fresh MRS, were measured at 600 nm. Surfaceome proteins from planktonic cells (1.5 mL) after overnight incubation were harvested by centrifugation (4000×*g*, 3 min, 4 °C). Cells were washed once with ice cold 100 mM sodium acetate buffer (pH 4.7). Cells from biofilm cultures after 48 h of incubation were harvested by removing non-adherent cells and washing adherent cells with ice cold acetate buffer as above. Cells normalized to equal cell densities were centrifugated (4000×*g*, 3 min, 4 °C) and then suspended gently in two-fold concentrated Laemmli buffer (pH 7.0). Cells were incubated in Laemmli buffer on ice for 15 min and then an additional five minutes at RT. Supernatants containing the released surface-associated proteins were recovered by centrifugation (4000×*g*, 3 min, 4 °C). Equal volumes from each sample were subjected to 12% TGX™ Gel (BioRad) electrophoresis (1-DE) using 1x Tris-glycine-SDS as the running buffer. After 1-DE, proteins were transferred onto a PVDF membrane using the TransBlot Turbo™ Transfer System (BioRad) according to the manufacturer’s instructions. The membrane was then probed first with antibodies (1:4000) detecting surface-associated factors of GG (28) and then with IRDye® 800CW goat anti-Rabbit IgG (LI-Cor® Biosciences) (1:20000). After probing, the membrane was blocked using Odyssey Blocking buffer and washed with PBS (phosphate buffered saline, pH 7.4) according to the instructions provided by LI-Cor® Biosciences. The cross-reacting antigens were detected and quantified using an Odyssey® infrared imaging system (LI-Cor® Biosciences) and the AlphaView Alpha View 3.1.1.0 software (ProteinSimple, San Jose, CA, USA), respectively. The experiment was repeated three times (each with two technical replicates).

### Mucus adhesion assay

Adhesion to mucus was carried out as described previously (29, 30). GG was grown in MRS-G and MRS-F with H^3^-thymidine for metabolic labeling in planktonic and biofilm forms as described above in section 2.4 (1-DE immunoblotting). Briefly, the planktonic and biofilm cells were first washed with a low pH buffer (100 mM sodium acetate, pH 4.7), which prevented the release of adhesive moonlighting proteins (cytoplasmic proteins) from the cell surfaces. Then, the washed cells were allowed to bind the porcine mucin immobilized onto microtiter plate wells. After washing the cells to remove non-adherent cells, the radioactivity of the adherent cells was measured by liquid scintillation, and the percent of bacterial adhesion was determined by calculating the ratio between the radioactivity of the adherent bacteria and that of the added bacteria. The experiment was repeated twice with four to five technical replicates. An unequal variance t-test was used to determine significant differences between selected samples.

### Indirect ELISA for detecting changes in SpaC abundance

Planktonic and biofilm GG cells were grown on MRS-G and MRS-F and washed with sodium acetate buffer as above. Washed cells at the same density were treated with 10 μM 4,6’-diamidino-2-phenylindole (DAPI; Molecular Probes) to stain all bacterial populations and with antiserum raised against His_6-_SpaC to detect SpaC expression. Antiserum against purified His_6-_SpaC of *L. rhamnosus* GG was raised in rabbits using routine immunization procedures as described previously (31). The staining was carried out by using indirect immunofluorescence as described previously (32) with minor modifications. Briefly, GG cells were collected by centrifugation, washed once with 100 mM Na-acetate buffer (pH 4.7) and fixed with 4% (wt/vol) paraformaldehyde in phosphate buffered saline (PBS; pH 4.0) prior to detection with anti-His_6_-SpaC primary antibody and Alexa-488 (Invitrogen)-conjugated goat anti-rabbit IgG (1 μg/ml) secondary antibody and DAPI counterstain. After staining, the optical density (OD_600_) of the cells was adjusted to 1.0 with washing buffer, and the intensity of the fluorescence in the samples was measured using the Victor3 1420 multilabel counter (PerkinElmer). The experiment was repeated twice, each with five technical replicates. An unequal variance t-test was used to determine significant differences between selected samples.

### Cell-surface shaving and LC-MS/MS analyses

Surfaceome analyses were conducted with GG cells cultured in planktonic (24 h) and biofilm states (48 h) in MRS-G and MRS-F at 37 °C under microaerophilic conditions. Biofilms were formed using flat-bottomed 24-well plates with 2.5 mL of the indicated media containing suspended GG cells (OD_600_ = 0.1) per well as described above. After 48 h of incubation, non-adherent cells were discarded, and the adherent cells were washed with ice-cold 100 mM Tris-HCL (pH 6.8). Pelleted cells suspended gently in 100 μl of TEAB were mixed with 50 ng/µL of sequencing grade modified porcine trypsin (Promega), and digestions were incubated at 37 °C for 15 min. Released peptides and trypsin were recovered by filtration through a 0.2 µm pore size acetate membrane by centrifugation (7000×*g*, 2 min, +4 °C), and the digestions were further incubated for 16 h at 37 °C. Digestions were stopped by adding trifluoroacetate (TFA) to a final concentration of 0.6%. The peptide concentrations were measured using a Nano-Drop (ND 1000, Fisher Scientific) at 280 nm. For each growth mode, three biological replicate cultures were used.

Trypsin-digested peptides were purified using ZipTips (C18) (Millipore), and equal amounts of the peptide samples were subjected to LC-MS/MS using an Ultimate 3000 nano-LC (Dionex) and QSTAR Elite hybrid quadrupole TOF mass spectrometer (Applied Biosystems/MDS Sciex) with nano-ESI ionization as previously described (28). The MS/MS data were searched against the concatenated GG protein database composed of the target (Acc. No. FM179322; 2944 entries) (12) and the decoy protein sequences using the Mascot (Matrix Science, version 2.4.0) search engine through the ProteinPilot software (version 4.0.8085). The Mascot search criteria were based on trypsin digestion with one allowed mis-cleavage, fixed modification of carbamidomethyl modification of cysteine, variable modification of oxidation of methionine, 50 ppm peptide mass tolerance, 0.2 Da MS/MS fragment tolerance and 1+, 2+ and 3+ peptide charges. The false discovery rate (FDR) percentages calculated using the formula 2xn*reverse*/ (n*reverse* + n*forward*) (33) indicated FDRs < 4% in each data set.

The raw Mascot proteomic identifications and associated files are freely downloadable from the the Dryad Digital Repository ().

### Protein identifications and surfaceome comparisons

Mass spectra from each replica sample were searched both separately and combined against the indicated protein database. Proteins with Mascot score [ms] ≥ 30 and p < 0.05 and identified in at least 2/3 replicates were considered high quality identifications and used to indicate condition-specific identifications and relative protein abundance changes. The emPAI (<u>e</u>xponentially <u>m</u>odified <u>P</u>rotein <u>A</u>bundance <u>I</u>ndex) values of the high confidence identifications were used to estimate protein abundance changes; emPAI is considered roughly proportional to the logarithm of the absolute protein concentration, which allows label-free and relative quantitation of the protein pairs (34). Principle component analysis (PCA) of the emPAI values was performed with IBM SPSS Statistics v.24. Missing values were substituted with half minimum emPAI value for each protein identified in at least 2/3 replicates in at least one of the conditions using MetImp 1.2 (https://metabolomics.cc.hawaii.edu/software/MetImp/) (35, 36) **(Table S1)**. Two proteins with zero variance (YP_003172193.1; YP_003172584.1) were excluded. PCA of the imputated emPAI data was carried out utilizing Varimax rotation with Kaiser normalization.

### Analysis of surface-associated aminopeptidase activities

Cell surface-associated aminopeptidase activity (PepN and PepC) was determined from planktonic and biofilm cells cultured in the presence of 2.0% and 0.5% glc and frc for 24 h (planktonic and biofilm cells) and 48 h (biofilm cells) using a previously reported method (37) with the following modifications. Briefly, biofilm and planktonic cells (1.8 mL) obtained from 10 mL cultures were washed as described above and then suspended in 1 mL of cold 100 mM Tris-HCl buffer, pH 6.8. The optical density of each sample was measured at 540 nm. The incubation mixture contained 1 mM L-leucine-*para*-nitroanilide (Leu-*p*NA) substrate in 200 µl of the cell suspension. The inhibitory effect of the metal-ion chelator EDTA was examined by incubating the cell suspension with 5 mM EDTA for a few minutes prior to the addition of Leu-pNA. Reactions were incubated at 37 °C for 5 min – 25 min and stopped with 800 µl of 30% (v/v) acetic acid. After centrifugation (8000×*g*, 5 min), the absorbance was measured at 410 nm. The specific aminopeptidase activity was calculated by dividing the absorbance change at 410 nm by the reaction time used for Leu-*p*NA hydrolysis and the optical density of the cells (OD_540_). An unequal variance t-test was used to determine significant differences between selected samples.

## Results

### Biofilm formation of GG on fructose and glucose

Biofilm formation of GG was first tested at three time points (24, 48 and 72 hours after inoculation, hpi) in MRS-G and MRS-F (2% glc and 2% frc) and MRS without carbohydrate, which indicated that frc stimulated biofilm formation by approximately 2-fold compared to glc at each time point tested (**Figure 1A**). The most efficient biofilm growth with frc and glc was detected at 48 and 72 hpi, respectively. The cell viability analysis of biofilm cells grown on glass-bottom dishes with both carbon sources revealed that biofilms formed on frc at 48 hpi contained proportionally more living cells than those grown on glc (**Figure 1B**). In addition, the biofilm formation efficiency was higher on hydrophilic polystyrene than hydrophilic glass under the conditions used (data not shown).

**Figure 1.**
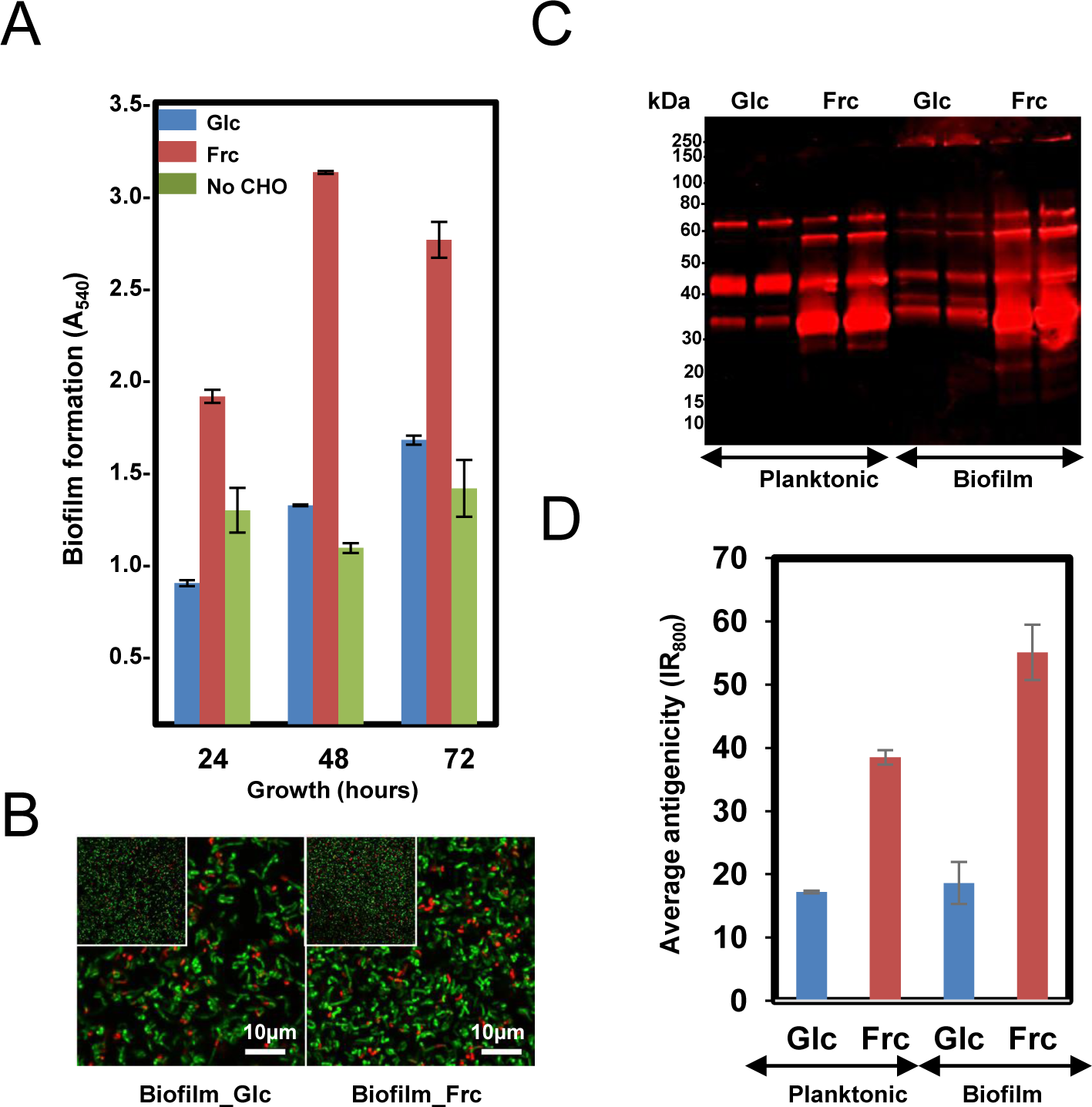
Assessing the biofilm formation of the GG cells in MRS in the presence 2% glc, 2% frc or without carbohydrate (CHO) under anaerobic conditions (5% CO_2_). **A)** Biofilm formation efficiency of the GG cells in the presence of frc and glc on flat-bottom polystyrene wells at indicated time points of incubation. **B)** Fluorescence images of 48 h old biofilms prepared in the presence of glc and frc. Biofilms were stained with the LIVE/DEAD *Bac*Light kit with Syto9 for staining viable cells and PI for dead cells. The scale bar is 10 μm. **(C)** 1-DE immunoblot analysis of planktonic and biofilm GG cells using anti-GG antibodies targeting the surface-associated proteins. The surface proteins were isolated from 24 h old (planktonic) and 48-h-old (biofilms) cells, and the amounts of samples separated by 1-DE were normalized to cell density. The antigen profiles were detected using an Odyssey® infrared imaging system. **(D)** Total fluorescence of each 1-DE antigen profile was quantified using the AlphaImager gel documentation and image analysis system.

### Growth mode- and carbon source-induced changes in cell-surface antigenicity

The surface antigenicity of GG cells grown in biofilm and planktonic states in MRS-G and MRS-F was studied by subjecting surface-associated proteins extracted in 1x Laemmli buffer to 1-DE combined with immunoblotting using antisera raised against intact GG cells. **Figure 1C** reveals that in each sample, the most intense/abundant cross-reacting antigen signals migrated to approximately 65, 55, 40 and 35 kDa. Comparing the total antigen intensity profiles, reflecting antigen abundances (**Figure 1C, lower panel**), showed that the presence of frc in the growth medium increased the surface antigenicity by ∼two-fold in planktonic and ∼ 2.5-fold in biofilm cells compared to cells grown on glc. In glc-associated cells, switching from planktonic to biofilm growth had no effect on surface-antigenicity, whereas the surface-antigenicity increased by ∼20% after switching from planktonic to biofilm growth in the presence of frc. The presence of Frc in the growth medium resulted in the appearance of unique protein bands at 15 and 20 kDa in both the planktonic and biofilm cell samples, while a higher-molecular weight protein (∼130 kDa) was specific to glc- and frc-biofilm samples. Thus, growing cells in planktonic or biofilm forms on frc increases the overall surface-antigenicity of GG along with the specific production of small-molecular-weight antigens, while a larger protein was specifically produced by biofilm cells growing on both carbon sources.

### Identifying the surfaceomes of GG cultured under different conditions

We previously demonstrated that the biofilm formation of GG is mainly protein-mediated (19). To explore this further, the biofilm-associated surfaceomes were identified by LC-MS/MS and identified proteins (p < 0.05) are listed in **Table S2**. Four surfaceome catalogs ([ms] ≥ 30, p < 0.05) were generated with each data set showing extensive overlap; 70% - 94% of the identified proteins were shared between at least two of the replica samples **(Table S3)**. The total number of high-quality identifications was 74, 106, 175 and 218 proteins from the glc-planktonic, frc-planktonic, glc-biofilm and frc-biofilm cells, respectively **(Table S4)**. As no decrease in the colony forming units (CFUs) of the shaved cells was observed (data not shown), we concluded that no significant cell lysis had occurred under the trypsin-shaving conditions used. Thus, comparison of the surfaceome catalogs indicated that the number of proteins at the cell surface of GG increased with frc as the carbon source compared to glc. Similarly, growth in the biofilm state increased the number of protein identifications compared to cells grown in planktonic form, regardless of the carbon source used.

### Multivariate analysis for screening specific surfaceome patterns

A principal component analysis (PCA) was performed to identify carbon source- and growth mode-dependent patterns in the surfaceome data. **Figure 2A** shows that all biological replicate samples clustered together well, with four clearly separated and identifiable groups. PC1, correlating with the change in growth mode-, and PC2 with the carbohydrate-dependent changes, suggest that the carbon source had a greater effect on GG during biofilm growth than planktonic growth. This was further explored by Venn diagrams that compared the number of specifically identified proteins (unique to growth mode and/or carbon source) and proteins with emPAI values showing ≥ 2-fold change **(Table S4)**. Venn digrams in **Figure 2B** compares the growth-mode and carbon source-associated surfaceomes. The number of specifically identified proteins (not identified from planktonic cells) was 106 and 115 in biofilm cells grown on glc and frc, respectively. In total, 6 and 3 identifications were specific for planktonic cells grown on glc and frc, respectively. In addition, the abundance of 37 and 58 proteins increased (≥ 2-fold) in biofilm cells, whereas 1 and 3 proteins were more abundant (≥ 2-fold) in planktonic cells with glc and frc, respectively. With frc as the carbon source, the number of specific identifications (not identified from glc-grown cells) from the planktonic and biofilm cells was 39 and 56, respectively. For glc, 7 and 13 specific identifications were made from the planktonic and biofilm cells, respectively. Two and 12 proteins displayed ≥ 2-times higher abundances in planktonic cells grown of glc and frc, respectively. The number of more abundant proteins with ≥ 2-fold change was 13 and 64 on glc- and frc-associated biofilms. Altogether, the presence of frc in the growth medium resulted in a higher number of specifically identified proteins as well as proteins with increased abundances in comparison to cells grown on glc.

**Figure 2.**
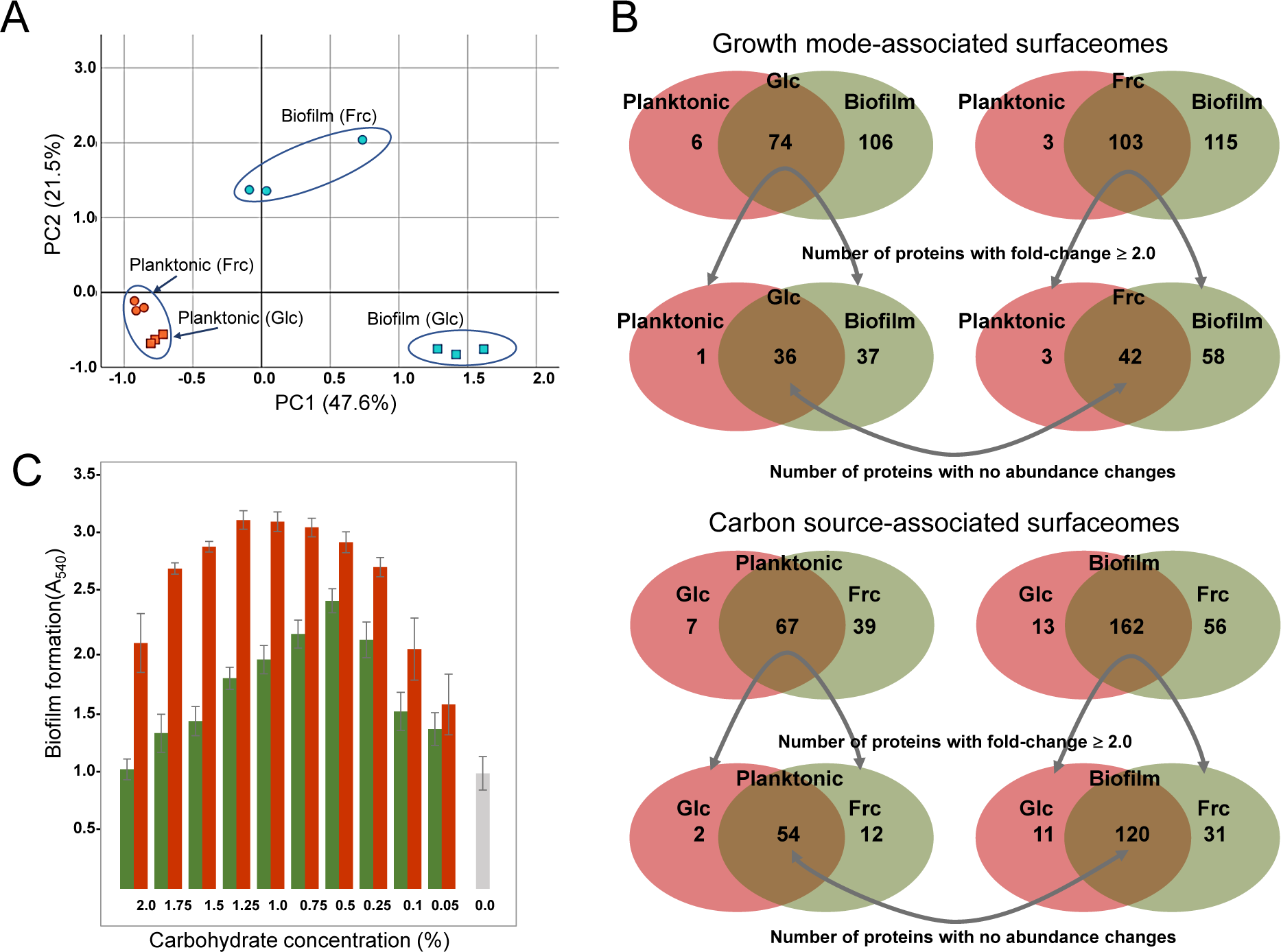
**(A)** Surfaceome variability among GG cells cultured under four different conditions (two carbon sources and two growth modes) assessed by PCA. PC1 (growth mode) accounts for 47.6% and PC2 (carbon source) accounts for 21.5% of the variation. **(B)** Venn diagrams indicate the number of commonly and specifically identified surfaceome proteins. **(C)** Biofilm formation of GG in varying concentrations of fructose (red bars) and glucose (green bars) and with no carbohydrate (grey bar). The biofilm formation efficiency was monitored after 48 h of incubation at 37 °C in the presence of 5% CO_2_.

Because Venn diagrams implied that the carbon source affected the number and amount of proteins attached to the biofilm surfaces, we also explored the biofilm formation efficiency in the presence of varying carbohydrate concentrations. **Figure 2C** shows that frc enhanced the biofilm formation over a wider concentration rate compared to glc. However, 0.5% glc produced the thickest biofilms at 48 hpi, while over two-fold higher concentrations of frc were required to achieve the thickest biofilms under the same conditions. Furthermore, optical densities (at 600 nm) of the formed biofilms grown on different carbohydrate concentrations revealed only marginal changes in cell densities, indicating that increased biofilm formation was not accompanied by an increased number of cells (data not shown).

### Identifying specific changes in protein abundance

The identification data were next compared to screen for individual growth mode- and carbon source-induced changes in the cell surface. The most distinctive protein abundance variations, including condition-specific identifications and proteins with abundance change ≥ 2-fold between the indicated conditions are listed in **Table S5**. The abundance changes are visually illustrated by heatmaps; **Figure 3** shows relative abundance changes estimated for classical surface proteins and known/adhesive moonlighters and **Figure 4** new or novel moonlighting proteins. Altogether, these proteins could be categorized into three groups: **(i)** the classically secreted surface adhesins, transporters and enzymes, **(ii)** known surface-associated moonlighters with adherence features, and **(iii)** new or novel moonlighters lacking established extracellular function. The known and predicted moonlighters were the dominant protein group among the detected surfaceomes. Among the identified moonlighters, the ribosomal proteins r-proteins (42 proteins), PTS/ABC-type transporter proteins (17 proteins; several OppA paralogs) and amino acid tRNA synthetases (10 proteins) formed the largest protein groups.

**Figure 3.**
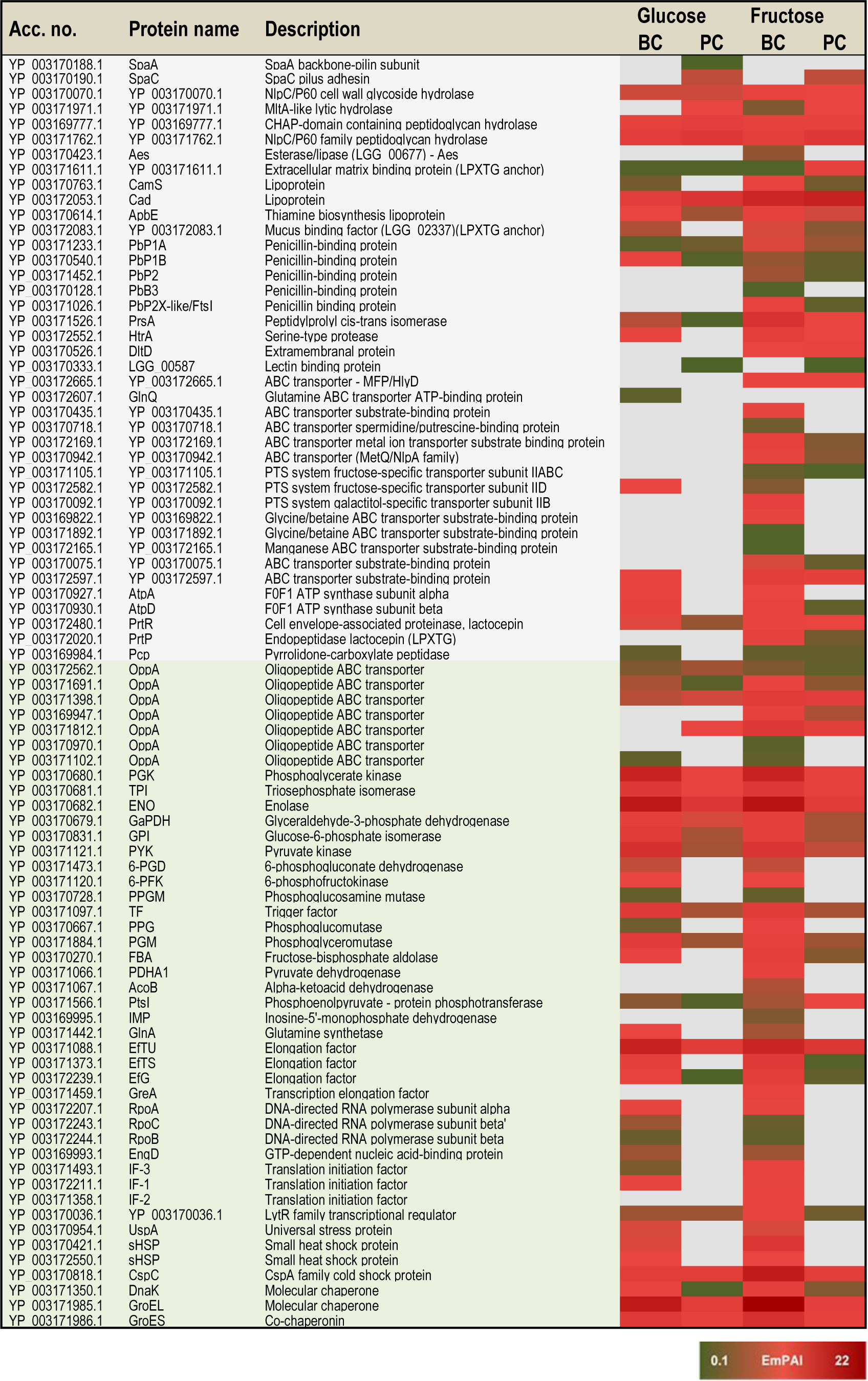
A heat map comparing the most distinctive protein abundance changes (estimated by the emPAI values) among the classical surface proteins anchored to cell-wall/-membrane via motifs or domains (names shaded in grey) and among known and/or adhesive moonlighters (names shaded in light green). Red and green refer to higher and lower protein abundances, respectively.

**Figure 4.**
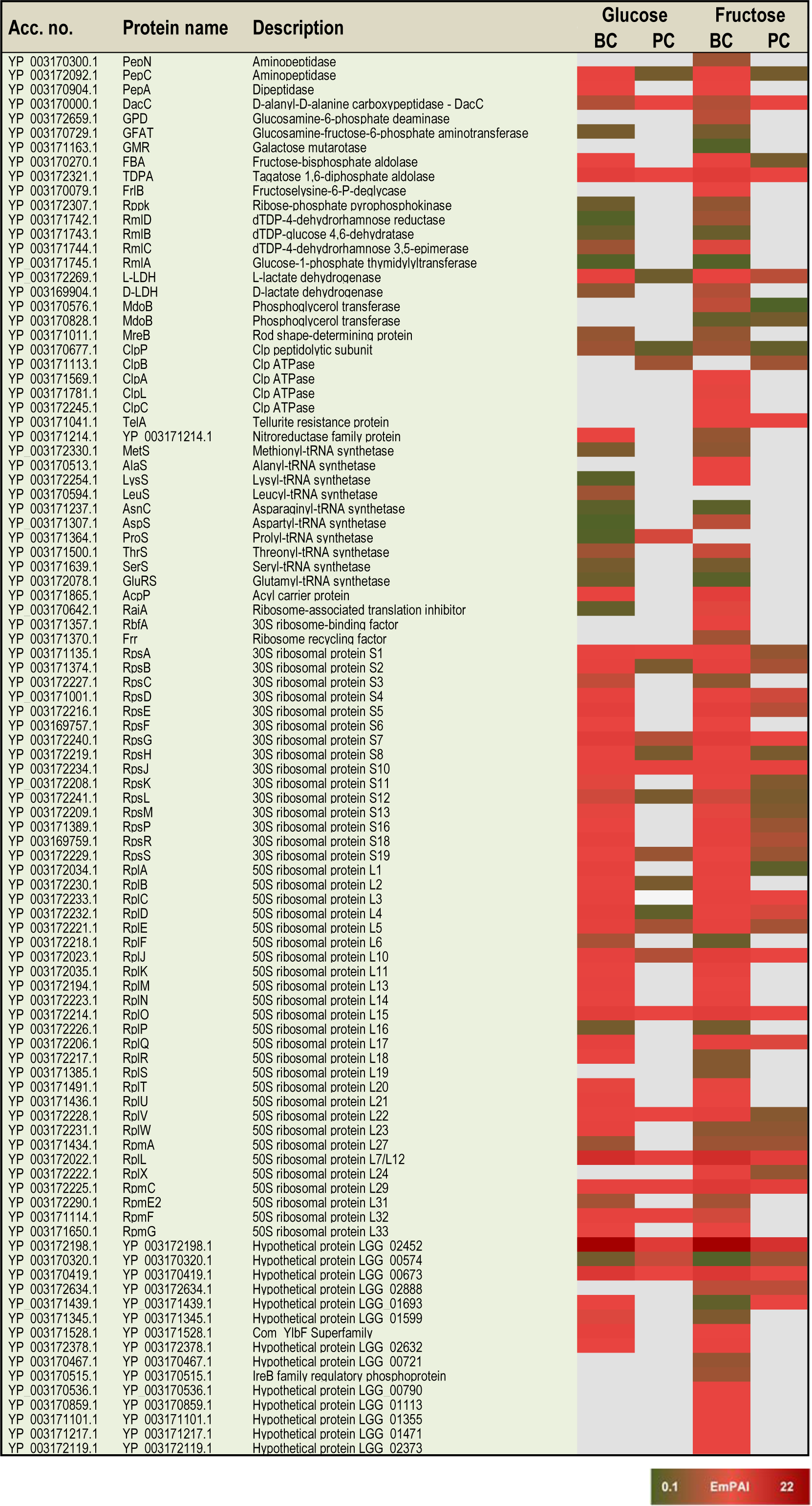
A heat map comparing the most distinctive protein abundance changes (estimated by the emPAI values) among the predicted moonlighters. Red and green refer to higher and lower protein abundances, respectively.

#### Proteins specific to one of the tested conditions

The greatest proportion (35 proteins) of the one-condition identifications was detected from the frc-biofilm cells. These included known and predicted moonlighters (e.g., RbfA, PDHA, GreA, GPD, FrlB, IF-2 and AlaS), Clp family chaperones (ClpL, ClpA and ClpC) and a PepN aminopeptidase, which were all proposed to be present at reasonably high abundances. Seven small molecular weight (6-33 kDa) moonlighters (YP_003170092.1, YP_003170859.1, YP_003170536.1, YP_003171217.1, YP_003171101.1, YP_003172119.1 and YP_003170467.1) were also found to be specific to frc-biofilm surfaces. The classical surface proteins, such as a 70-kDa penicillin-binding protein (PbB3) and a 28-kDa Aes lipase, were specific to frc-biofilm surfaces and, based on the emPAI values were predicted to be produced at low levels under these conditions. Leucyl-tRNA synthetase (LeuS) and glutamine transport (GlnQ) were specifically identified from glc-biofilm surfaces. Protein identifications specific to planktonic cells growing on glc included the shaft pilin subunit SpaA, encoded by the *spaCBA* pilin operon (12).

#### Growth mode-specific identifications

In total, 3 and 47 proteins were detected as specific to planktonic and biofilm cell growth, respectively, independent of the carbon source. Planktonic-specific identifications included the classically secreted 97-kDa SpaC pilin subunit (encoded by the *spaCBA* operon), a 73-kDa lectin-binding protein and the moonlighting Clp family ATPase ClpB. Classical surface-proteins specific to biofilm cells included one of the OppA transporter paralogs (YP_003171102.1) components of the F-ATPase (F1Fo ATPase) membrane bound complex, AtpA confering catalytic and beta AtpD regulatory functions. AtpA was specific to biofilm cells growing on glc and frc. AltD was more abundant than AtpA on biofilm cells, while the beta subunit was also detected in planktonic cells grown on frc. Several known and predicted moonlighters were also specifically identified from biofilm cell surfaces. These included seven amino acid tRNA synthetases (Thr, Met, Asn, Ser, Glu, Lys and Asp); 11 known (D-LDH, PPGM, 6-PGD, UspA, IF-1, 6-PFK, RpoB/C, GlnA, PepA, and EngD); and 13 predicted moonlighters (RpsF, RplP, RpsQ, RpmE2, RplT, RpmG, RplM, RpsC, RplF, RplR, RmlA, RmlC and RmlD). Among the predicted moonlighters, RmlA, RmlC and RmlD are encoded by the four-gene (*rmlABCD*) operon involved in dTDP-rhamnose biosynthesis.

#### Carbon source-specific identifications

From the two carbon sources, frc was found to induce specific surfaceome changes similar in both the planktonic and biofilm cell surfaces, whereas only one glc-specific identification (prolyl-tRNA synthetase-ProS) was shared by the planktonic and biofilm cells. The frc-specific identifications included classical surface proteins, such as the lactocepin (PrtP), an extramembranal protein (DltD), a tellurite resistance protein (TelA), five ABC/PTS-type transporters mediating type I protein secretion (MFP/HlyD), amino acid (OppA)/metal ion intake/output proteins (YP_003172665.1, YP_003169947.1, YP_003169947.1, YP_003170942.1, YP_003172169.1, YP_003170075.1 and YP_003171105.) and two penicillin-binding proteins (PbP2 and PbP2X-like/FtsI). In addition, two phosphoglycerol transferase (MdoB) paralogs of different sizes (78 and 84 kDa) and an r-protein, RplX, were detected as moonlighting proteins specific to frc-associated cells. From these, the greatest differences were associated with RplX, the ABC transporters for metal ion binding (YP_003172169.1) and oligopeptide uptake (OppA, YP_003169947.1), the 78-kDa MdoB paralog and the penicillin-binding proteins PbP2X-like and PbP2, which all displayed over two-fold higher abundances during biofilm growth compared to planktonic growth on frc.

#### Growth mode-specific changes

Planktonic- or biofilm growth-induced surfaceome changes independent of the carbon source used were considered growth mode-induced changes. Twenty-nine proteins with similar abundance levels in both the glc- and frc-biofilm cells and higher abundances in comparison to their planktonic counterpart cells included several known and predicted moonlighters. Among these, the greatest differences in abundance were detected for predicted moonlighters, such as an elongation factor – EfG (>8.0-fold increase), an 8-kDa hypothetic protein – YP_003172198.1 (with ∼6.0-fold increase), an aminopeptidase – PepC (with ∼5.0-fold increase) and a ClpP caseinolytic peptidase (>2.0-fold increase). Known moonlighters (PGK, TPI, L-LDH, GaPDH and RpoA) and 21 r-proteins (RplR, RplO, RpsG, RplJ, RplU, RplQ, RpsD, RpsE, RpsR, RpsD, RpsE, RpsB, RpsS, RpsA, RpsM, RpsH, RplK, RpsL, RplN, RplK and RplB) were less abundant but still over 2-fold more abundant on biofilm than on planktonic cell surfaces. Only one of the OppA paralogs (YP_003171102.1) was found to be ∼5-times more abundant on planktonic cells on both carbon sources.

#### Carbon source-induced changes in biofilm cultures

In total, 61 proteins more abundant in glc and frc biofilms than in planktonic cells were divided into four groups as follows: **(i)** Proteins “more induced” on glc than on frc, i.e., proteins exhibiting greater relative abundance difference between planktonic and biofilm growth modes when grown on glc. This group included twenty-two proteins and 14 proteins with > 5-fold higher differences in abundance and included the classically secreted penicillin-binding protein (PbP1B) as well as several known and predicted moonlighters (DnaK, L-LDH, PYK, PGM, GBI, RplA, RplD, RpsE, RpsB, RplQ, RpsG, and the 8-kDa YP_003172198.1). **(ii)** Proteins more abundant on glc-biofilm than on frc-biofilm cell surfaces included dihydrolipoamide acetyltransferase (∼4.7-fold increase), a potential Com_YlbF family protein predicted to ensure proper biofilm formation (∼2.7-fold increase) and five r-proteins (RplK, RplB, RplN, RpsF and RpmF) with fold-changes ranging from 2.4 – 4.6. **(iii)** Proteins “more induced” on frc than on glc. The majority of these were identified as known and predicted moonlighters (EfTS, GroEL, AtpD, FBA, RplV, RpsR and CspC), with fold-change 4-15 and with EfTS (∼15-fold increase) and GroEL (∼9-fold) displaying the greatest changes in abundance. The classical surface-proteins, including the PrtP protease (specific to frc-cells) and the lipoprotein CamS (∼4-fold increase), were predicted to be more abundant on frc cells than on glc cells. **(iv)** Proteins more abundantly produced on frc-than on glc-associated biofilm cell surfaces included five classical surface proteins, which included one of the OppA paralogs (YP_003171812.1), the PrsA chaperone/foldase, the serine-proteases PrtP and HtrA and the CamS lipoprotein. Of these, OppA (>15-fold increase), PrsA (∼7-fold increase) and PrtP (∼4-fold increase) displayed the greatest changes in abundance. In total, 9 known and predicted moonlighters were also more abundantly produced (≥ 4-fold) on frc-biofilm cell surfaces, including an 18 kDa small heat shock protein (sHSP), a cold-shock protein (CspC), two amino acid tRNA synthetases (LysS and AspS), RpsR, phosphoglucomutase (PPG), and a dTDP-glucose-4,6-dehydratase (RmlB, encoded by the *rmlABCD* operon). In addition, a dTDP-4-dehydrorhamnose reductase – RmlD, with a >2.0-fold change in abundance, was also identified from the frc-biofilm cells.

#### Carbon source-induced changes in planktonic cultures

Carbon source-dependent changes in planktonic cell surfaces could be divided into two groups. **(i)** Proteins predicted to be more abundant on cells growing on frc included the extracellular matrix-binding protein (YP_003171611.1) with >8-fold higher abundance. Another classical surface protein, PrsA, and two moonlighters (PtsI and RplC) were also predicted to be ∼5-fold more abundantly produced on frc cells. **(ii)** Proteins predicted to be more abundantly produced in the presence of glc included one of the OppA paralogs (YP_003171812.1), an MltA lytic hydrolase, a prolyl-tRNA synthetase – ProS, and the AtpD protein.

### Adherence of biofilm and planktonic cells to porcine mucus

Because the identification data implied that the growth mode and carbon source in the growth medium could affect the adherence features of GG, we investigated the mucus-binding ability of the planktonic and biofilm cells grown on glc and frc. **Figure 5A** indicates that the biofilm cells grown on frc displayed the highest level of adherence, while planktonic cells grown on glc were the least adherent. The frc biofilms demonstrated a mucus-binding ability that was ∼2-fold more efficient than that of the glc-associated planktonic cells; p = 0.037). The possible role of the SpaC-adhesin, the tip pilin of the SpaCBA pilus known to mediate key interactions with the human mucus (12), in mediating the frc-stimulated adherence was assessed next by testing the adherence of the GG cells in the presence of anti-SpaC antiserum. **Figure 5A** indicates that the presence of SpaC antibodies markedly decreased the adherence of each type of cell to mucus. Although the final level of adherence detected for these cells was somewhat similar, the biofilm cells grown on frc displayed the greatest SpaC-mediated inhibition, whereas the level of adherence decreased the least in glc-associated planktonic cells. To readdress the possible role of the pilus structure in the observed differences, we also monitored the SpaC abundances on each GG cell type. As shown in **Figure 5B**, the highest abundance of SpaC was detected on the glc-associated biofilm cells, which was >2-fold (p = 4.46e-09) higher than that on the planktonic cells grown on glc and ∼2-fold (p = 1.15e-07) higher than that on the frc-biofilm cells. Planktonic cells grown on frc displayed the lowest SpaC abundance.

**Figure 5.**
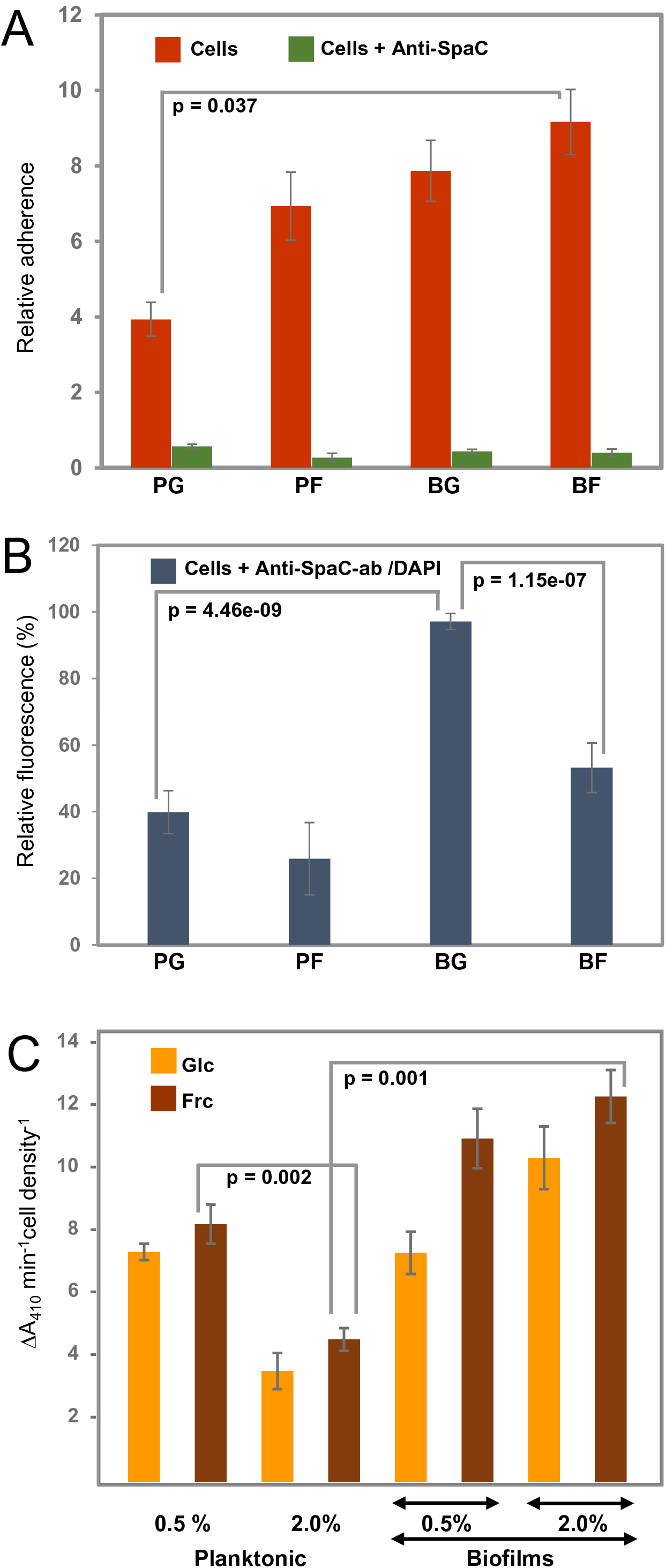
Representative diagrams **(A)** showing the relative adherences (average adhesion percentage of the added bacteria) of planktonic (PC) and biofilm GG (BC) cells grown on frc and glc to porcine mucus and **(B)** illustrating the abundance of the SpaC pilin subunit on the cells treated first with anti-SpaC-antibody and then Alexa-488-conjugated goat anti-rabbit IgG and/or the DAPI counterstain. **(C)** The surface-associated aminopeptidase activity (PepC and PepN) of the planktonic and biofilm cells was assessed using 1 mM Leu-pNA. Significant differences are indicated with lines. The error bars indicate SD for two or more biological replicates, each with several technical repeats.

### Whole-cell aminopeptidase activity

Varying abundances detected for predicted cell-surface moonlighters, such as PepN, PepC, PepA, ClpP, and several OppA paralogs, implied differences in the ability of the GG cells to utilize casein/casein-derived peptides for growth. PepN was specific to frc-biofilm cells. PepC was predicted to be equally abundant on the biofilm cells grown on frc and glc and displayed over a 4-fold higher abundance on biofilm than planktonic cells. Comparison of PepN and PepC implies that PepC is produced ∼2-fold more on the frc-biofilm cells than PepN on the same cells. To verify these results, we monitored the surface-associated aminopeptidase activity by exposing washed and intact GG cells to a chromogenic substrate, Leu-*p*NA, a specific substrate for both aminopeptidases (37). **Figure 5C** confirms the identification findings by showing that the surface-associated aminopeptidase activity of frc-biofilm cells was ∼4-fold higher (p = 0.001) than that of planktonic cells growing on frc. Changes from glc to frc increased the aminopeptidase activity on both the planktonic and biofilm cell surfaces but led to a proportionally greater increase in the biofilm cell surface. Lowering the carbohydrate concentration to 0.5% increased the aminopeptidase activity in planktonic cells by ∼4-fold, while only a slight decrease in the enzyme activity was observed in biofilm cells on both carbon sources. Enzyme activities were also examined in the presence of 5 mM EDTA, a metal-chelator that inhibits the activity of metallopeptidases such as PepN while not affecting other types of peptidases such as PepC (37). Since EDTA inhibited only marginally the total aminopeptidase activity only in frc-biofilm cells, PepC is the likely candidate for the hydrolysis of the substrate.

## Discussion

### Carbon source controls the protein-dependent biofilm growth of GG

Environmental signals, such as O_2_, CO_2_, bile, mucins, nondigestible polysaccharides and carbohydrates present in the human GIT, have been shown to affect the GG biofilm formation *in vitro* (15, 19). In MRS growth medium, GG produces biofilm, which is completely disintegrated by treatment with proteolytic enzyme (19), indicating the presence of a large amount of proteins in the biofilm matrix and/or that biofilm formation is protein-mediated. Since both carbon metabolism (15) and cell-surface exposed proteins (19) appear to play a role in biofilm formation, we explored this link further by investigating the effects of glc and frc as carbon sources on the surfaceome composition of GG. Our findings indicated that changing the carbon source from glc to frc resulted in enhanced biofilm formation efficiency of GG *in vitro*. GG is known to exert high *in vitro* adhesion to Intestinal Epithelial Cells (IECs) and biofilm formation capacity on polystyrene and glass (15, 38). Here, our findings demonstrated more efficient biofilm growth on polystyrene than on glass and more efficient biofilm growth with frc than with glc on both inert materials. Crystal violet provides a good detection of biofilm mass, but this dye stains both the bacterial cells and extracellular matrix, including proteins (39, 40). As the tested carbohydrates had only a marginal effect on the cell density, we suggest that the increase in crystal violet staining, instead of showing increased biofilm formation, related directly to changes in the protein abundance of biofilm cells. A similar observation was also made when growing the biofilm cells with varying carbohydrate concentrations; decreased carbohydrate concentrations resulted in thicker biofilms, which did not result from higher cell density (data not shown). Thus, changes in carbon sources and their concentrations regulate the changes in protein abundance on the biofilm cell surfaces.

### Biofilm growth and frc in growth medium enhance the protein export in GG

Examining the cause-effect relationships between the identified surfaceomes revealed that the carbon source played a more important role in the biofilm mode of growth. Conditions of glc limitation have been shown to enhance the biofilm formation of GG (16). Here, decreasing the concentration of both carbon sources in growth medium resulted in similar outcomes. The presence of frc enhanced biofilm formation over wider concentrations than glc. Known and novel moonlighting proteins were the dominant protein group in the biofilm-associated surfaceomes. Comparing the frc- and glc-associated surfaceomes implied that the export of these nonclassical surface proteins is more efficient in both the biofilm and planktonic cells growing on frc. The presence of moonlighting proteins on the cell surface or in the culture medium has been reported for several microbial species, and for many of these, the moonlighting activity outside of the cells has been demonstrated (41, 42). It has been proposed that bacteria recycle conserved cytoplasmic proteins in a pH-dependent manner in the matrix to facilitate interspecies interactions without specifically recognizing the dedicated matrix components of the other species (43). In a recent study, the strongly positively charged r-proteins were shown to be embedded in *S. aureus* biofilm matrix under acidic conditions [44]. This was proposed to depend on the pH, coordinated by the formation/release of acidic fermentation end-products in the biofilm cells facing oxygen limitation [44]. In the present study, the pHs of the spent culture supernatants were clearly acidic, pH < 5.0 (data not shown). Thus, while r-proteins were identified as the most abundant moonlighters in the biofilm matrices, we suggest that low pH could have promoted the interactions between the r-proteins and the biofilm cells also in the present study. Although mechanisms that signals cytoplasmic proteins out from the cells are not known, a WXG motif present in some moonlighters or the SecA2-dependent secretion system are considered potential signals/mechanisms driving the export of nonclassical moonlighters.

In general, the mechanisms underlying their export/transport remain unknown (45). In view of this, it is tempting to speculate that the specific appearance of HlyD/MFP in frc-associated cell surfaces indicates the role of this protein in directing moonlighting proteins out of the cell. HlyD is part of a translocon that comprises HlyB, HlyD and TolC, which is proposed to coordinate the transport of potential novel proteins by quite different mechanisms (46). Overexpression of PrsA, a known surface-associated chaperone or foldase acting on secreted proteins (47), on biofilm cells and frc-associated cells is another plausible factor contributing to protein secretion under these conditions. This chaperone/foldase is known to enhance protein secretion efficiency in bacteria (48) and act on various substrates, ranging from 20 to 80 kDa in size (49). The serine-type surface-protease, HtrA, follows the same abundance trend as PrsA and is another classical surface-protein that could enhance protein secretion in frc-associated samples. In bacteria, HtrA is involved in degrading abnormal proteins, processing secreted proproteins and the maturation of native proteins (50), but this protease can also affect biofilm formation and control the presence of surface-associated moonlighters, such as enolase (ENO) and glyceraldehyde-3-phosphate dehydrogenease (GaPDH) (51).

Here, ENO, EfTU, TDPA, PGK and GroEL, each with previously reported adhesive functions (41, 42), were detected as the most abundant moonlighters in the biofilm cells growing on both carbon sources. In addition, stress response proteins (GroES and DnaK), glycolytic proteins (PGM, PYK, PGK, FBA, GPI, L/D-LDH, GMR and PDCE2), the elongation factors EfTS and EfG, the RNA polymerase subunit RpoA, the translation initiation factor IF1, and the trigger factor TF, were also detected as relatively abundant on biofilms. The r-protein moonlighters (a total of 43 proteins) were the largest protein group among the biofilm-associated surfaceomes, independent of the carbon source used. In addition to their proposed role as biofilm integrity/stability enhancing role, coordinated by the production of acidic fermentation end-products (44), some of the identified r-proteins may also have other moonlighting functions. The most abundant r-proteins were RplX, RpsE, RpsG, RplL and RplO, from which RpIL (the 50S ribosomal protein L7/L12) was predicted to be the most abundant. The high abundance of this r-protein during biofilm growth could explain the concomitant presence of another moonlighting protein, EfG, on the biofilm cells at high abundances, as the GTPase activity of EfG is reported to require RplL for ribosomal translocation (52). For several r-proteins (e.g., L5, L11, L23, L13a, S3, S19, S27), noncanonical functions ranging from gene expression regulation to subverting pro-inflammatory actions, apoptosis and protection against abiotic stress (41, 53, 54) have been proposed. The increased abundance of the trigger factor (TF) during the biofilm mode of growth could be explained by the concomitant presence of the ribosomal protein L23. This r-protein has been reported to have binding sites for both the TF and the signal recognition particle, which is thought to aid in protein export to the cell membrane (55). The specific appearance of several moonlighting aminoacyl-tRNA synthetases (ATRSs) in biofilm cells was also interesting, as these enzymes, unrelated to their primary function, also regulated gene expression, signaling, transduction, cell migration, tumorigenesis, angiogenesis, and/or inflammation (41, 56), thereby forming a novel group of moonlighters in probiotic species.

### Fructose and biofilm growth increase the surface-associated aminopeptidase activity in GG

Several components of the proteolytic system (several OppA paralogs, PepC, PepN and PepA) involved in the utilization of milk casein (19) were identified here as novel moonlighters with higher abundances on biofilm than planktonic cell surfaces. Among these PepN and the OppA paralogs were more abundant on or specific to the frc-associated cell surfaces. The enzymatic assay performed on whole cells using a substrate specific to PepC and PepN confirmed the identification results and indicated that more PepC than PepN is associated with biofilm cells. In addition, the classical surface proteases PrtP and PrtR were predicted to be more abundantly produced on the biofilm cell surfaces. The two identified aminopeptidases are expressed in the cytoplasm, where they act on the oligopeptides that are taken up by the Opp-transport system. Thus, efficient surface-aminopeptidase activity could provide the probiotic with means to speed up growth and adaptation in conditions involving oligopeptides as the carbon source. In a recent study by Galli et al. (57) GG was utilized as an adjunct starter culture in Camembert-type cheese production, which was found to improve the sensory characteristics of the cheese. While the mechanism behind the GG-mediated flavor formation in cheese remains to be elucidated, the efficient and diversified proteolytic system of the strain could be involved (57). It is highly likely that the robust GG cells do not undergo autolysis during cheese making, which makes the cell-surface located moonlighting aminopeptidases C and N interesting objects to be studied further in this context.

Several cytoplasmic proteins have been shown to be selectively sorted into membrane vesicles (MVs), which in some bacteria were shown to contribute to approximately 20% of the whole biofilm matrix proteome, demonstrating that MVs are also important constituents of the biofilm matrices (58). Considering this finding, we suggest that among the classical and nonclassical proteins, PepA, PepN, PepC, the RNA polymerase subunits RpoA and RpoB, an Aes lipase and stress-chaperone GroEL could be targeted to MVs, which is an important mechanism exploited by both gram-negative and gram-positive bacteria to export proteins in a protected and concentrated manner to aid neighboring cells or modulate the host immune system. Taken together, our findings suggest that the export of many classical and nonclassical proteins could be enhanced in GG when cells are grown in a biofilm state.

### Surface-antigenicity and adherence of GG by biofilm growth in the presence of frc

Comparison of the total antigenicity profiles of GG cells grown in planktonic and biofilm forms on glc and frc indicated that frc as the carbon source increases the abundances of many antigens (∼23, 28, 40, 60 and 80 kDa in size) during both growth modes. We previously demonstrated the presence of several adhesive and antigenic moonlighters at the cell surface of planktonic GG cells grown on glc (28). In that study, protein moonlighters, such as DnaK (67.2 kDa), GroEL (57.4 kDa), PYK (62.8 kDa), TF (49.8 kDa), ENO (47.1 kDa), GaPDH (36.7 kDa), L-LDH (35.5 kDa), EfTu (43.6 kDa), Adk (23.7 kDa) and UspA (16.8 kDa), were identified as highly abundant and antigenic on planktonic GG cells. The present study suggests that PYK, ENO, GaPDH, TF, L-LDH, EfTU, UspA, DnaK and GroEL could be produced more during biofilm than planktonic growth, which is also in line with an earlier proteomic study showing that a switch from planktonic to biofilm growth in *Lactobacillus plantarum* increases the abundances of several stress responses and glycolytic moonlighters (59). From the identified moonlighters, ENO, IMP, FBA, TDPA, GreA, GroES and GroEL are plausible factors that aid in biofilm formation in the presence of frc, as evidenced by surfaceome predictions.

The *rmlABCD* operon products, involved in the synthesis of O-antigen lipopolysaccharide (60), could have conferred increased surface antigenicity to GG cells during biofilm growth. Here, all of the operon gene products RmlA, RmlB, RmlC and RmlD with glucose-1-phosphate thymidylyltransferase, dTDP-4-dehydrorhamnose 3,5-epimerase, dTDP-4-dehydrorhamnose reductase and dTDP-glucose-4,6-dehydratase activities, respectively, were detected only in biofilm cells and with ∼2-fold higher abundances in the presence of frc. In some pathogens, L-rhamnose is required for virulence, and the enzymes of the Rml pathway are considered potential targets in drug design (61). In view of this, detection of the Rml enzymes as moonlighters is interesting and points towards a novel role of these enzymes at the cell surface of nonpathogenic bacteria. The rhamnose moiety has also been shown to bind specific moonlighters (e.g., ENO and 58 other potential moonlighters) at the bacterial cell surface, proposing that rhamnose-mediated anchoring is a general mechanism for anchoring the moonlighting proteins to the cell surface in bacteria (62).

The specific appearance of MdoB, a phosphoglycerol transferase involved in the biosynthesis of lipoteichoic acid (LTA), could also be considered a potential antigen in GG. Inactivating the *mdoB* gene in a probiotic *Lactobacillus acidophilus* (strain NCK2025) was shown to protect mice against induced colitis (63). Here, two MdoB paralogs of different sizes were increasingly produced in frc biofilms and planktonic cells, and their predicted MWs (78 and 84 kDa) could explain the increased abundance of an antigen protein band in frc samples migrating to approximately 80 kDa. In relation to MdoB, the identified DltD, encoded by part of the *dltABCD* operon involved in the formation of lipoteichoic acid (LTA), could have increased antigenicity in frc-associated GG cells. DltD is a single-pass membrane protein involved in the final transfer of D-alanine residues to LTA on the outside of the cell (64) that contributes to adherence and biofilm formation (65, 66). LTA has been shown to be essential for biofilm matrix assembly and bulk accumulation over time (67), and the early stages of biofilm formation necessitates efficient expression of *dltD* (68). In addition, a lack of DltD has been linked to poor acid survival in planktonic cells and an inability to form biofilms *in vitro* in the presence of sucrose or glucose (69). As DltD could not be detected in the glc-associated planktonic or biofilm cells, we hypothesize that such deficiency might also explain the lower biofilm formation efficiency on glc compared to frc.

Many of the classical surface proteins may have also contributed to the total antigenicity and/or adherence in the GG cells. The most plausible candidates include lipase and lipoproteins (Aes, AbpE, Cad and CamS), peptidoglycan hydrolases (e.g., the NlpC/P60), mucus-binding factor (MBF), a lectin-binding protein, penicillin-binding proteins and two pilus proteins (SpaC and SpaA). From these, the SpaA backbone-pilin subunit, SpaC adhesin and MBF are most known; the first two are produced through the *spaCBA* operon coding for the pilus, the key factor promoting biofilm formation *in vitro* as well as colonization and adherence *in vivo* (15, 70), while MBF has been shown to bind intestinal and porcine colonic mucus, laminin, collagen IV, and fibronectin (71, 72). Additional mucus-adherence assays with each G cell type in the presence of anti-SpaC antibodies complemented with indirect ELISA monitoring of the SpaC abundances indicated the importance of the pilus adhesin in coordinating the adherence in both the planktonic and biofilm cells. The inability to identify SpaA or SpaC from biofilm-cells implied that the overwhelming presence of moonlighting proteins on biofilm cell surfaces accompanied by the complex structure of the pilus adhesin could be the reason why the two pilus proteins were identified only from the planktonic cells. The same reason applies for the lack of MabA among the identified surfaceomes. MabA (LGG_01865), the modulator of adhesion and biofilm formation, is proposed to strengthen the biofilm structure following the pilus-mediated interaction with the surface (16).

A recent study highlighted an important role of glc and frc, carbohydrates prevalent in the Western diet, in regulating the colonization of a beneficial microbe independent of supplying these carbohydrates to the intestinal microbiota (26). In that study, both frc and glc, but not prebiotics such as fructo-oligosaccharides, were found to silence the two-component system sensor histidine kinase/response regulator Roc that activates transcription of clustered polysaccharide utilization genes in a widely distributed gut commensal bacterium *B. thetaiotaomicron*. Roc was also suggested to promote gut colonization by interacting with moonlighting proteins such as GPI or glucose 6-phosphate dehydrogenase (73). From these two moonlighters, GPI was detected here as a more abundant protein in biofilm cells with a slight increase in cells grown on glc. As the present study compared surfaceomes of cells grown only in the presence of simple carbohydrates, we cannot exclude the possibility that corresponding response regulators (e.g., WalK) could have been modulated by these carbohydrates and that the two tested carbohydrates could coordinate the colonization of GG *in vivo* in an analogous manner.

## Conclusions

The present study provides the first in-depth comparison of the planktonic- and biofilm matrix-associated surfaceomes of GG. We show that cells growing in planktonic and biofilm forms in the presence of simple carbohydrates in growth medium could be used to modify the surfaceome composition of this probiotic. We show remarkable differences among the compositions of the classical and nonclassical surface proteins (moonlighters) that have immunomodulatory, adherence, protein-folding, proteolytic or hydrolytic activities. Our study also indicated that the carbon source coordinates protein-mediated biofilm formation, as evidenced by a whole-cell enzymatic assay measuring the surface-associated aminopeptidase activity on cells cultured on different carbon sources and with varying carbohydrate concentrations. The total antigenicity and adherence were higher on biofilm and planktonic cells grown on frc, in which specific surface-adhesins and/or known and novel moonlighters were the plausible contributory factors. Our findings also demonstrated the key role of the SpaCBA pilus independent of the growth mode or carbon source used, whereas the increased protein moonlighting is suggested to strengthen the biofilm structures and/or aid in cell-cell interactions. The observed phenotypic variations in *L. rhamnosus* GG potentially includes probiotic (adherence and immunomodulatory) and industrially relevant (proteolytic activity) features. Whether the GG phenotypes could be modulated in the host or in the food conditions remains to be shown.

## Funding

This study was supported by the Academy of Finland (Grant No. 272363 to PV and 285632 to VK) and by Sigrid Juselius Foundation’s Senior Reseacher’s grant to RS

## Conflict of Interest Statement

The authors declare that the research was conducted in the absence of any commercial or financial relationships that could be construed as a potential conflict of interest.

## Author Contributions

KS, PV, TN, PS, VK, RS, AF and JS conceived, designed, performed the experiments, IM analyzed the data, and KS and PV wrote the manuscript. All authors participated in the revision of the manuscript and approved the final version.

